# *ggDNAvis*: a *ggplot2*-based *R* package for creating high-quality DNA sequence and modification visualisations

**DOI:** 10.64898/2026.03.02.708895

**Authors:** Evelyn Jade, Emma L. Scotter

## Abstract

In the genomics age, enormous volumes of DNA sequencing information are continuously produced and analysed. Pipelines for processing DNA information are widespread and mature. However, tools for creating figure-ready visualisations of sequence and methylation information can be limited, often resulting in authors taking varied, manual approaches. *ggDNAvis* produces high-quality, extensively customisable renders of DNA sequences and modifications. The three core features are visualisation of a single sequence, multiple sequences, or methylation of multiple DNA molecules. Additionally, *ggDNAvis* has tools for reading/writing file formats such as FASTQ. Single-sequence visualisation accepts any DNA/RNA sequence from any source, so is extremely versatile and useful across biological contexts. Multiple-sequence visualisation was conceived in the context of visualising read-level long-read sequencing (e.g. Nanopore) daya to show intrasample variability in causative genes of repeat expansion diseases such as neuronal intranuclear inclusion disease (NIID) caused by *NOTCH2NLC*, Huntington’s disease (HD) caused by *HTT*, and fragile X syndrome (FXS) caused by *FMR1*, but is applicable for visualising any set of multiple sequences. Methylation visualisation requires read-level methylation data, most commonly obtained via modification-capable basecalling of signal-level Nanopore data. *ggDNAvis* is available as a *Shiny* web app and a CRAN *R* package, with the latter supporting additional *ggplot2* extension.

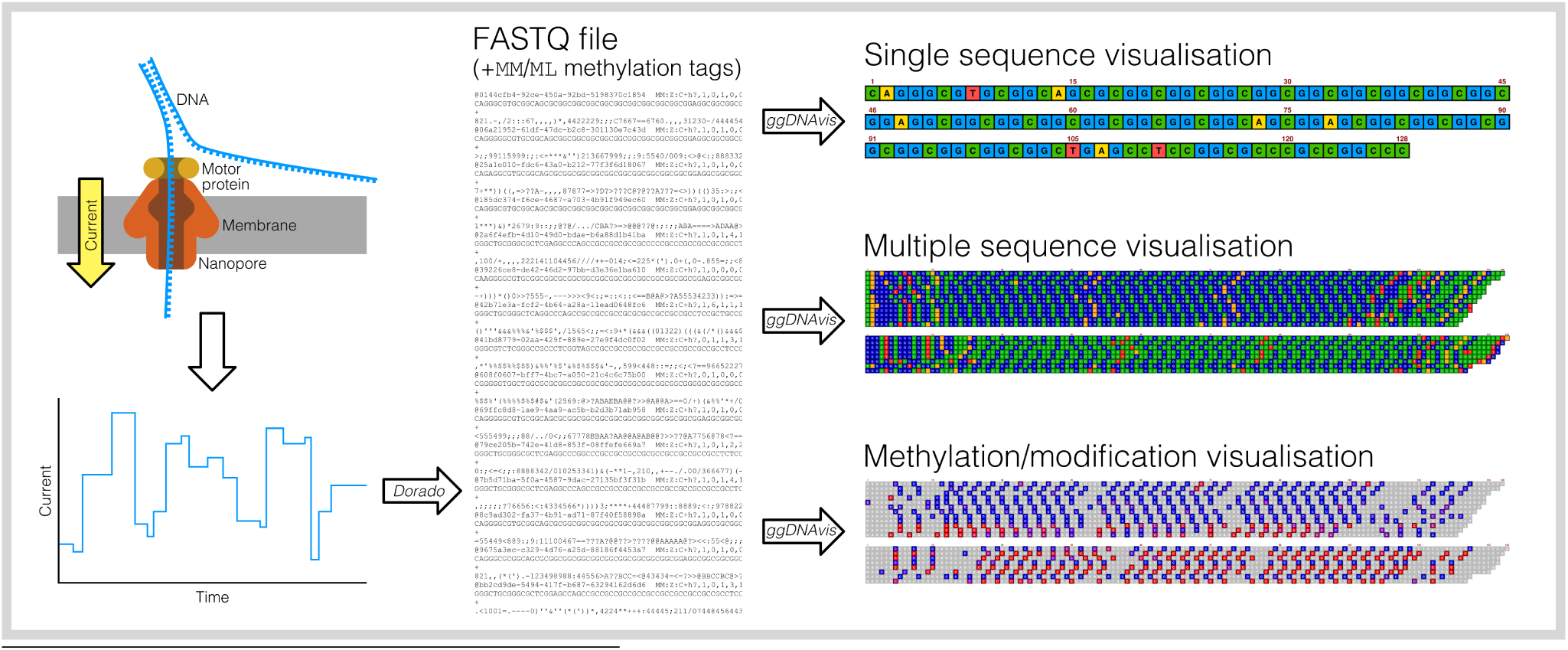

## 1 Background

Advancements in DNA sequencing technology have led to ever-increasing volumes of DNA sequencing data. The National Centre for Biotechnology Information estimated over 91 exabases (∼22,000 terabytes) were stored in the Sequence Read Archive database alone as of February 2024 (NCBI, 2024). There are well-established, near-universal standards for recording and processing DNA sequence information such as FASTA (Pearson & Lipman, 1988), FASTQ (Cock et al., 2010), and SAM/BAM (H. Li et al., 2009). However, at the end point of bioinformatic pipelines, visualisation of DNA outputs is unstandardised and often necessitates improvisation by authors. Some published approaches generate two-dimensional graphs to visualise the information content of a DNA string (Lee, 2019; T. Li et al., 2024) — however, there is no widely circulated tool to produce renders of the sequence itself. Other software suites such as *IGV: Integrative Genomics Viewer* (Robinson et al., 2011) and *Jalview* (Waterhouse et al., 2009) provide extensive analysis and functional viewing capabilities, but are not built to prioritise the creation of customisable, publication-quality sequence renders. The authors are not aware of any widely distributed software whose purpose is to create high-quality, highly customisable renders of DNA sequences.

One example of a field where the lack of such software is detrimental is the wide range of neurodevelopmental and neurodegenerative diseases caused by expansion of short tandem repeat (STR) regions, including conditions such as Huntington’s disease (HD) (caused by a CAG repeat expansion in the *HTT* gene) (MacDonald et al., 1993) and fragile X syndrome (FXS) (caused by a CGG repeat in the 5’ untranslated region (UTR) of the *FMR1* gene) (Verkerk et al., 1991). Neuronal intranuclear inclusion disease (NIID) is similarly a neurodegenerative disease that was linked in 2019 to a GGC/GGA repeat expansion in the 5’ UTR of the *NOTCH2NLC* gene (Deng et al., 2019; Ishiura et al., 2019; Sone et al., 2019; Tian et al., 2019) — this *NOTCH2NLC* work showcases a range of recent approaches taken to DNA visualisation in a practical medical genetics and bioinformatics context. *NOTCH2NLC* repeat expansions are frequently published as ‘consensus’ sequences summarising long-read sequencing information into a single representative sequence for each allele of each participant (Fitrah et al., 2023; Sone et al., 2019; Yu, Deng et al., 2021; Yu, Luan et al., 2021). However, presentation of these consensus sequences varies greatly in readability and accuracy across the *NOTCH2NLC* literature (**Figure 1**).

**Figure 1:**
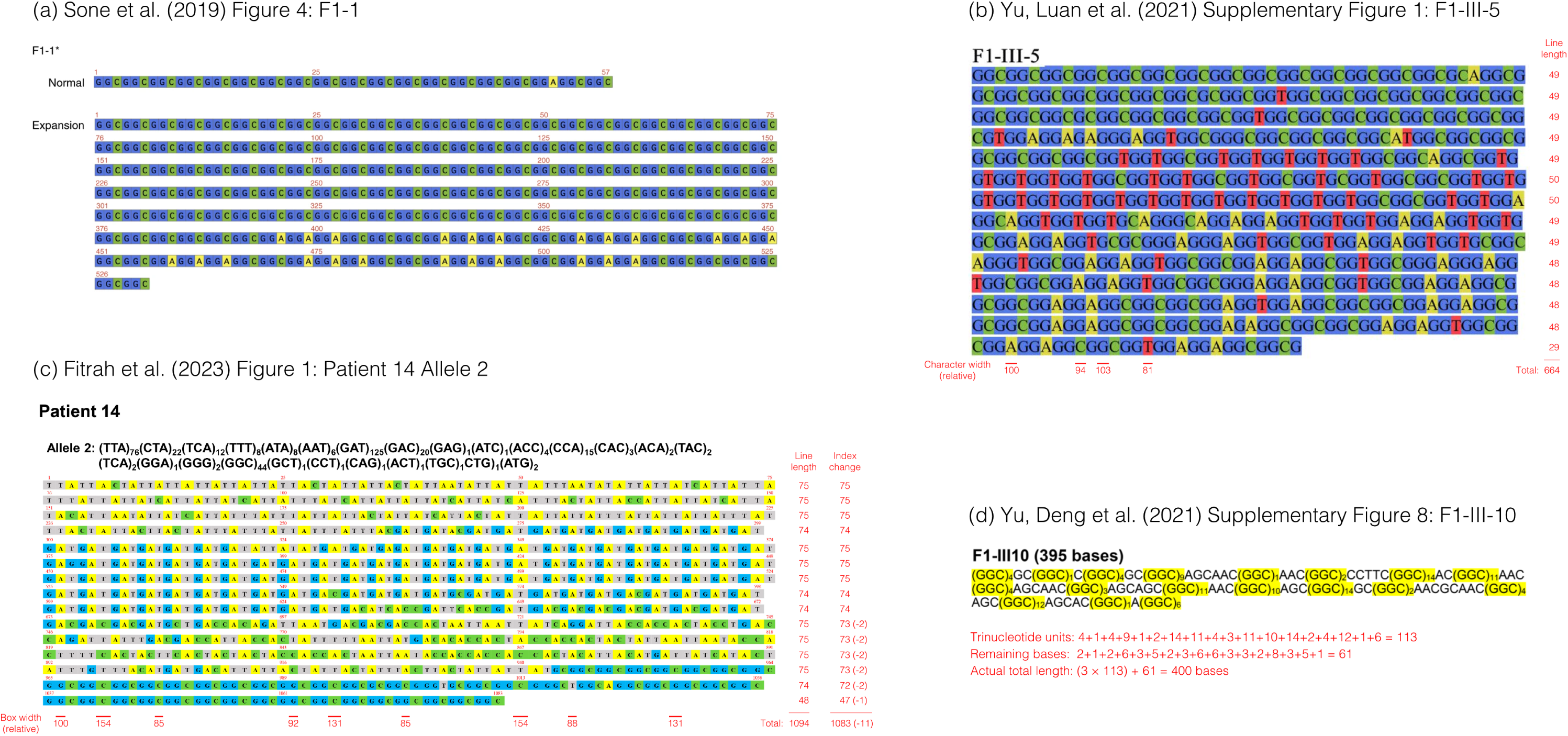
Different approaches to DNA visualisation across the *NOTCH2NLC* literature: **(a)** a uniform grid with identically sized and spaced boxes for each base, correctly numbered at a regular interval (Sone et al., 2019); **(b)** a screenshot of the sequence in a word processor, using a non-monospaced font that results in uneven character widths, inconsistent line ending positions, and variable numbers of bases per line (Yu, Luan et al., 2021); **(c)** a table with uneven line spacing and box widths, variable number of bases per line, and numerical errors in the index annotations (Fitrah et al., 2023); and **(d)** the sequence in a word processor with repeated GGC units condensed to (GGC)*_n_*, but with numerical error in the given total number of bases (Yu, Deng et al., 2021). Inconsistencies and numerical errors are annotated in red. Relative box/character widths were determined by matching the width of the base representation with a horizontal line in *Affinity*, then dividing the pixel width of each line by that of the leftmost line, multiplying by 100, and rounding. All visualisations are reproduced and annotated for the legally protected purpose of critique (fair use/fair dealing exception); scope of critique and comparison is specifically the DNA visualisation approaches and not any other scientific or biological features of the articles.

**Figure 2:**
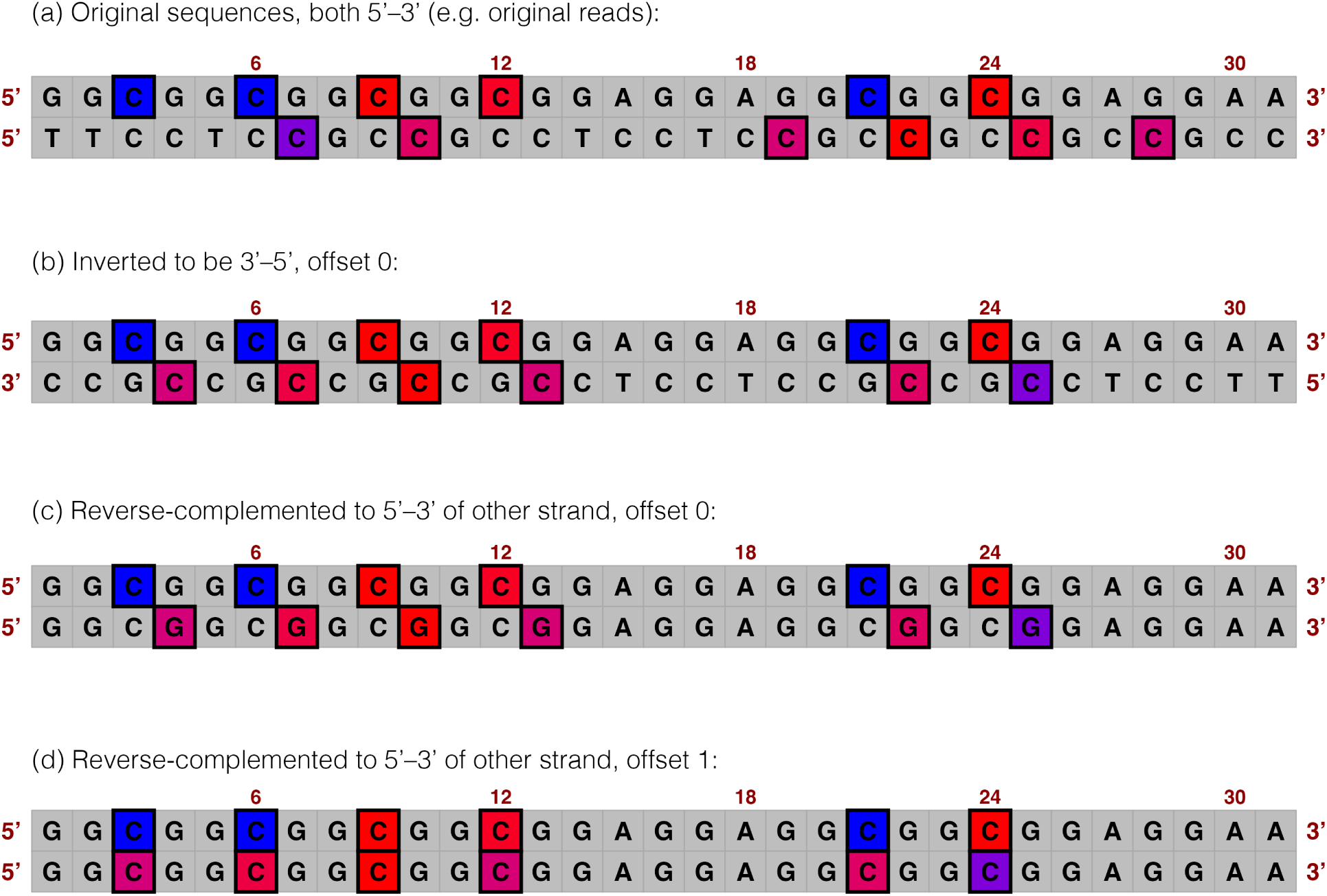
*ggDNAvis* handling of methylation of reverse reads. Two reads are shown over the same region but in opposite directions. The options for displaying reverse reads are **(a)** leaving reverse reads unchanged, **(b)** inverting to be complementary to forward reads, **(c)** reverse-complementing to be identical in sequence to forward reads, and **(d)** reverse-complementing and offsetting modification locations to be displayed on the C of CGs. All processing is done on the reverse (bottom) read; the forward (top) read is shown purely for comparison. Generated by location_reversing_example.R.

Selected examples include consensuses presented as regular grids coloured by base and annotated with index/base position numbers at regular intervals (**Figure 1a**; Sone et al., 2019), or, less successfully, a screenshot of plain (non-monospaced) text with black letters highlighted in colours corresponding to each base (**Figure 1b**; Yu, Luan et al., 2021). A third approach appears to approximate the grid style from **Figure 1a** but with inconsistencies in the size and spacing of the boxes that suggest it was created as a table in a word processor (**Figure 1c**; Fitrah et al., 2023). A final example shows the consensus sequence as text but with GGC repeat units condensed to (GGC)*_n_*and highlighted yellow (**Figure 1d**; Yu, Deng et al., 2021). In the latter two examples, errors in the annotated number of bases compared to the actual number in the given sequence (annotated red in **Figure 1**) suggest that the figures were created manually — this would be tedious and time-consuming for authors, and difficult for readers to interpret, but no easy automated solution appears to have been available. Of these varied approaches to DNA visualisation across the *NOTCH2NLC* literature, the organised grid style with index annotations at regular intervals (**Figure 1a**; Sone et al., 2019) was chosen for its consistency, reliability, and ease of reading. Therefore, emulating this style with an automated, easy-to-use, highly customisable tool was the principal motivation for creating *ggDNAvis*. *ggDNAvis* was conceived with the sole purpose of visualising DNA and exporting publication-ready renders. This enabled a tighter focus on render quality and customisability compared to comprehensive analysis suites such as *IGV* and *Jalview*. Built on the widely used *ggplot2* graphics system (Wickham, 2016) and returning *ggplot2* objects, *ggDNAvis* visualisations can also be further modified through standard *ggplot2* commands (e.g. **Figures 3–6**). Together, *ggDNAvis* allows visualisation of (a) single DNA sequences, (b) multiple DNA sequences, or (c) methylation of multiple DNA strands. There is also built-in capability for parsing the common FASTQ file format, including modification information (e.g. methylation) when present in header rows, and integrating metadata such as read directions and participant IDs from a CSV file. An interactive *Shiny* web application provides a graphical user interface (GUI) that removes *R* programming skills as a requirement for using *ggDNAvis*.

**Figure 3:**
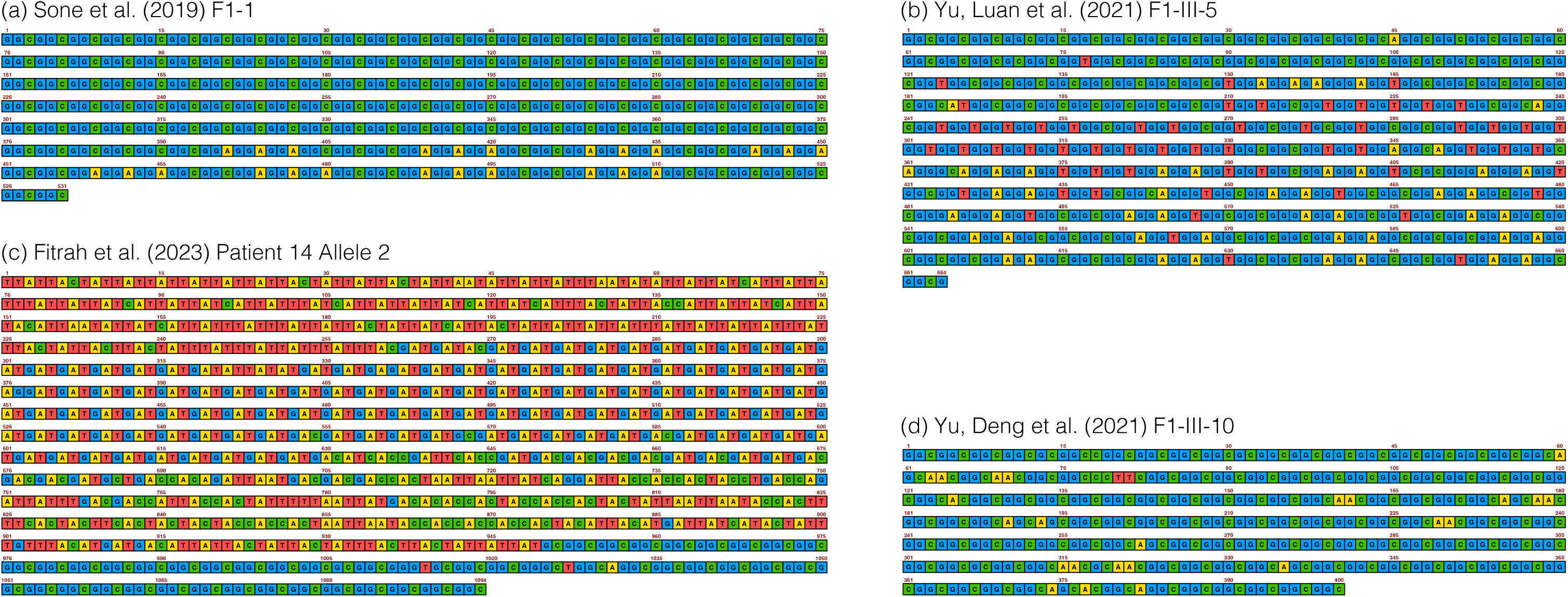
*NOTCH2NLC* consensus sequences from Figure 1 re-visualised using visualise_single_sequence(), including the expanded alleles of patients **(a)** F1-1 from Sone et al. (2019); **(b)** F1-III-5 from Yu, Luan et al. (2021); **(c)** Patient 14 (Allele 2) from Fitrah et al. (2023); and **(d)** F1-III-10 from Yu, Deng et al. (2021). Compared to the original visualisations reproduced in Figure 1, *ggDNAvis* visualisations should be significantly easier for authors to create because the process is highly automatic (while retaining customisability), and are also easier for audiences to read because they have a universal grid layout, fully consistent box sizes and line spacing, and no numerical errors in index annotation. Generated by single_sequence_ notch2nlc.R — note that while this code is fairly complex, the complexity comes from compiling multiple *ggDNAvis* visualisations into a single plot within *R*. Generating any one visualisation requires only a single function call, and compiling and labelling visualisations can easily be done in standard image editing software by users who so prefer.

**Figure 4:**
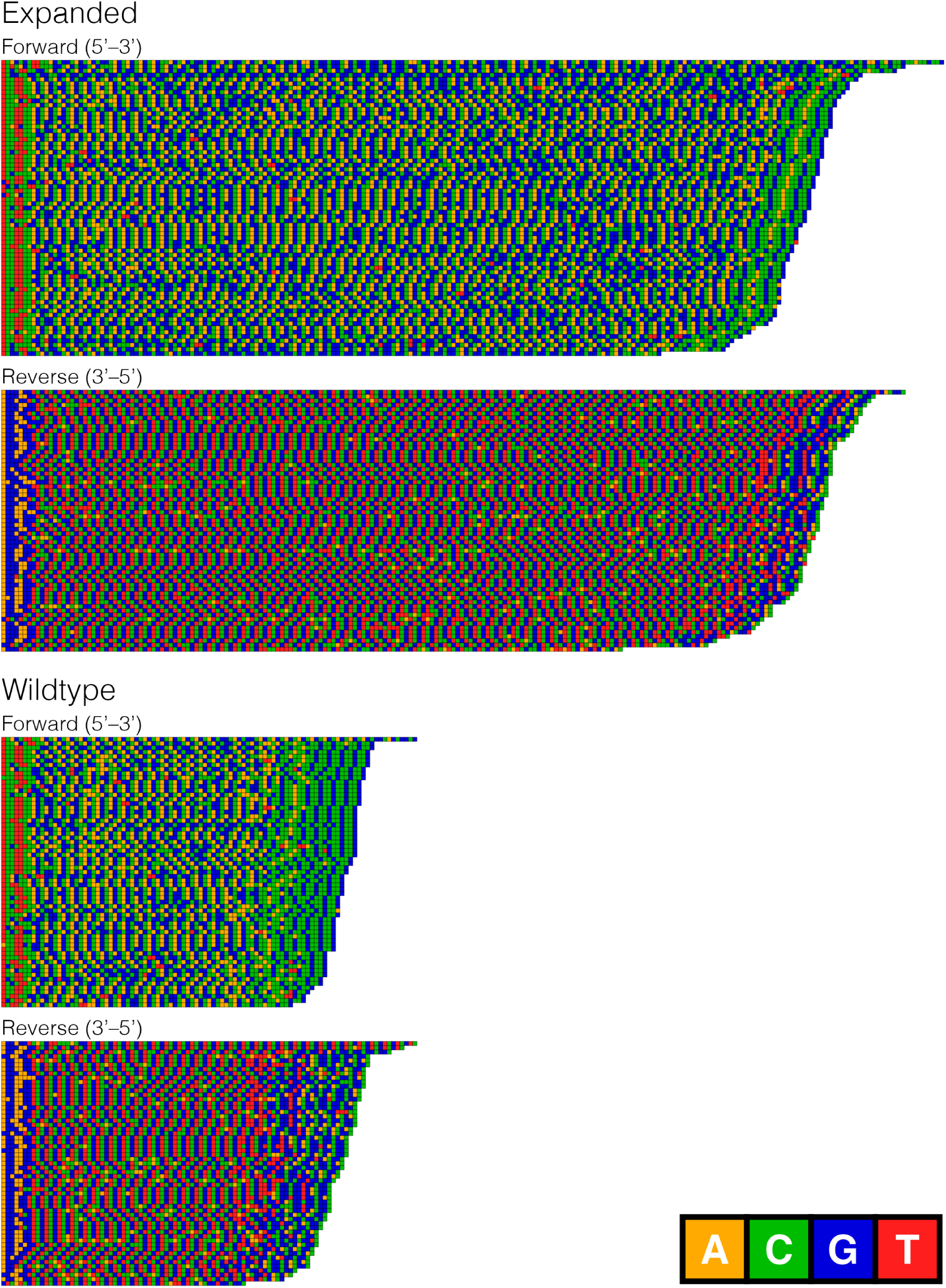
Clipped *HTT* reads from Nanopore DNA sequencing of HD fibroblasts visualised using visualise_many_sequences(). Separated by allele (expanded: length >138 bp; wildtype: clipped read length ≤138 bp) and read direction; reverse reads inverted to 3’–5’ to complement 5’–3’ forward reads. Sequencing data from Fang et al. (2022), downloaded from Sequence Read Archive (sample ND33392, accession SRR13068459, from study SRP292765). Reads were mapped, filtered, and clipped by many_sequences_htt.sh, then randomly subsampled to 250 reads and visualised in *ggDNAvis* by many_sequences_htt.R.

**Figure 5:**
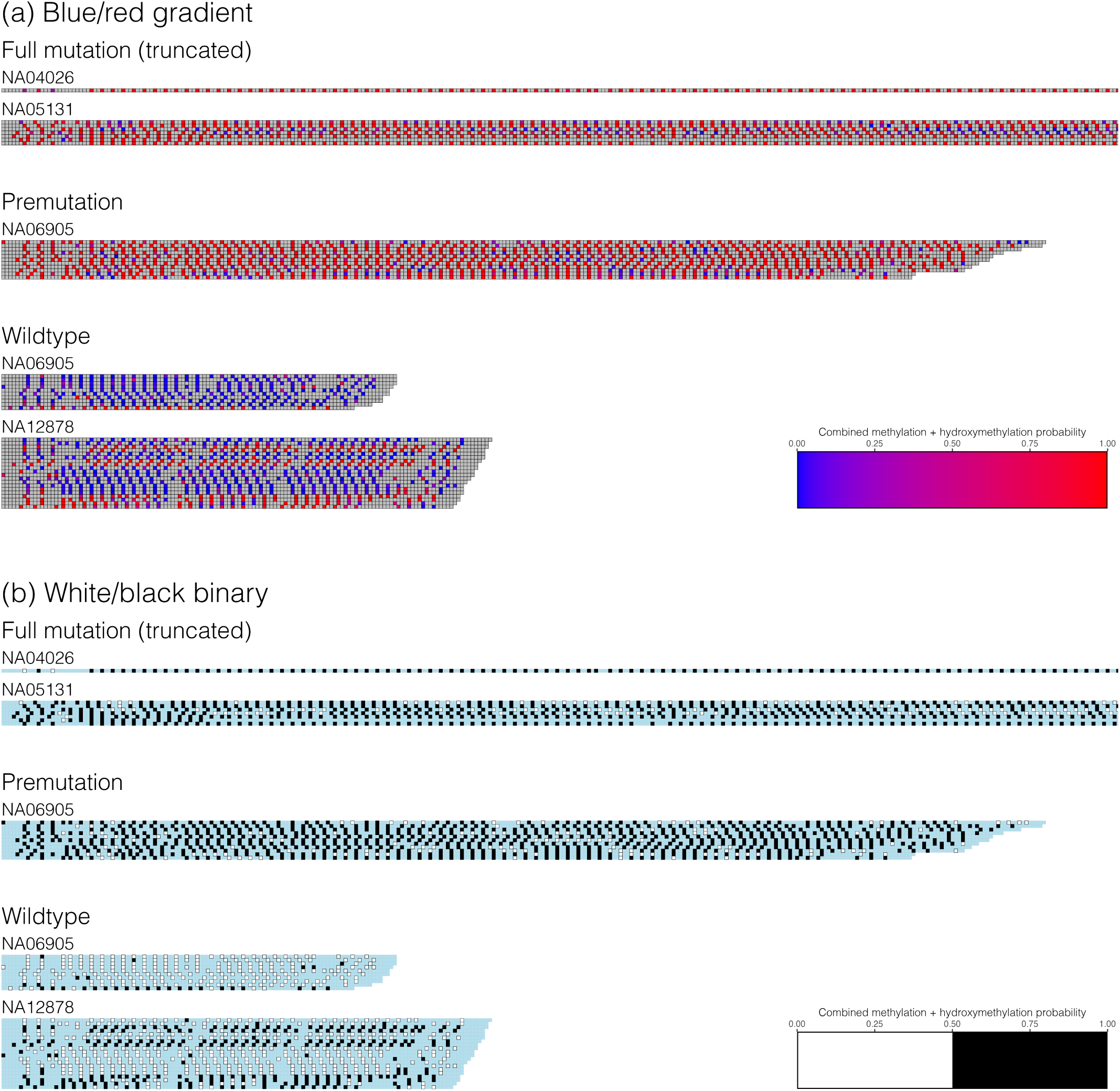
DNA cytosine modification of clipped *FMR1* reads from Nanopore sequencing of FXS affected, premutation carrier, and healthy control cell lines visualised using visualise_methylation(), with probabilities either. **(a)** shown in a blue/red gradient along the entire 0–1 probability range or **(b)** called as either unmodified (white, probability integer ≤127) or modified (black, probability integer ≥128). Modification was assessed for C bases in a CG context, with other bases (grey in **(a)**, light blue in **(b)**) not assessed for methylation. Modification probabilities are the combined scores of CG methylation (MM:C+m?) and hydroxymethylation (MM:C+h?). Sequencing performed by Stevanovski et al. (2022) on Coriell Institute cell lines NA04026 (FXS XY male, hemizygous full mutation), NA05131 (FXS XY male, hemizygous full mutation), NA06905 (unaffected XX female, heterozygous premutation), and NA12878 (unaffected XX female, homozygous wildtype). Data downloaded from SRA study SRP349335 via download_fmr1.sh, processed via methylation_ fmr1.sh, then figure generated by methylation_fmr1.R.

**Figure 6:**
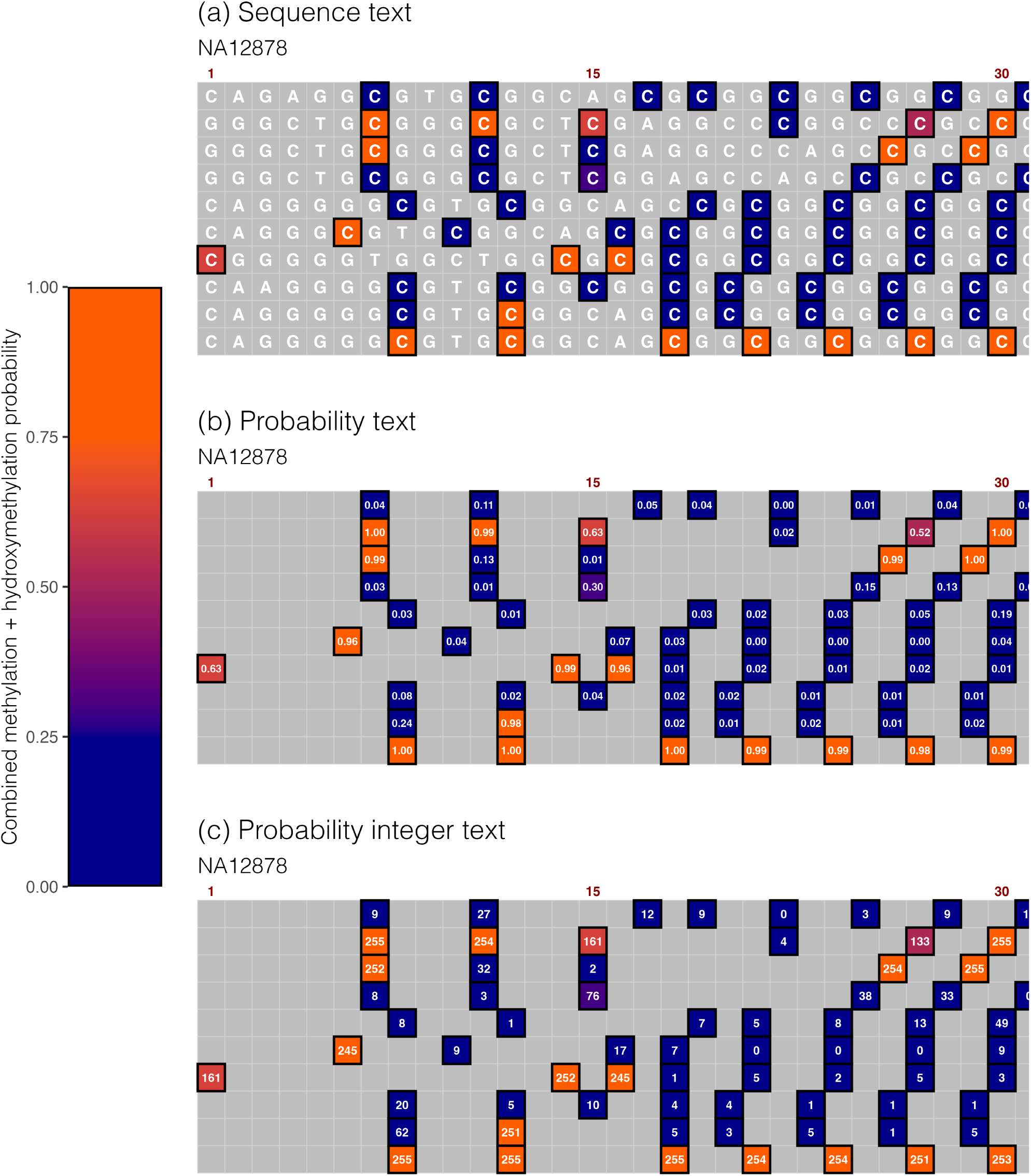
Text display options for DNA modification visualisation, demonstrated on truncated clipped *FMR1* reads from Nanopore sequencing of an unaffected XX female cell line visualised using visualise_methylation(). Options include **(a)** rendering the sequence letter inside every base, **(b)** rendering the 0–1 methylation probability inside each modification-assessed base (sequence_text_scaling = c(−0.5, 256), sequence_text_rounding = 2), and **(c)** rendering the 0–255 integer probability inside each modification-assessed base (sequence_text_scaling = c(0, 1), sequence_text_rounding = 0). Probability gradient clamped at 0.25*255 and 0.75*255 i.e. all modification probabilities ≤25% are uniformly blue, all modification probabilities ≥75% are uniformly orange, and probabilities from 25% to 75% are interpolated along the blue/orange gradient. Coriell Institute cell line NA12878 (homozygous wildtype) sequenced by Stevanovski et al. (2022), data downloaded from SRA study SRP349335 via download_fmr1.sh, processed via methylation_ fmr1.sh, then 10 reads randomly subsampled and figure generated by methylation_fmr1.R.

*ggDNAvis* is fully open-source under the permissive MIT licence. All code for the package itself, the accompanying *Shiny* app, generating the included datasets, and generating all coded figures in this paper is available from the source code repository. This manuscript discusses the software gap that prompted development of *ggDNAvis*, its implementation methods, and a summary of the types of outputs and customisation options available. A non-expert user guide (with examples) and full technical documentation for *ggDNAvis* is available from the accompanying documentation website.

## 2 Implementation

### 2.1 Data management

#### 2.1.1 FASTQ parsing

To enable provision of data to the *ggDNAvis* visualisation functions, FASTQ parsing functions are included within the package. The read_fastq() function reads FASTQ sequencing information to a dataframe by assigning each line of the file to read (unique ID), sequence, or quality columns based on line number modulo 4, where each row represents one read. Additionally, when base modification information is available for each read, such as from methylation-capable Nanopore long-read sequencing (Ambardar et al., 2016), BAM files can be written to FASTQ with modification information from the MM/ML tags stored in the FASTQ header rows via *SAMtools* (H. Li et al., 2009). The read_modified_ fastq() function can parse these ‘modified FASTQ’ files by splitting the header rows at tab characters to extract the read ID, the MM (modification location) tag, and the ML (modification probability) tag separately. Modification parsing is described in detail in the supplementary materal, but in summary the MM tag stores a list of which target bases (most frequently cytosine) were assessed for modification as the number of non-assessed bases in between each assessed base (Bonfield & Marshall, 2024). *ggDNAvis* converts this into a list of which indices along the sequence correspond to bases assessed for modification (e.g. 3,7 denotes the third and seventh bases). The ML tag stores the probability of each type of modification (e.g. methylation or hydroxymethylation) at each assessed base location as an integer from 0 to 255, where each integer *p* represents the probability space from 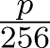 to 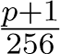 (Bonfield & Marshall, 2024). The integer probabilities are split by modification type and stored as a complementary list to the locations, so that the dataframe returned by `read_modified_fastq()` contains one column of modification locations and one of modification probabilities for each modification type, as well as a column listing all modification types present in each read.

#### 2.1.2 Metadata merging

To make a useful display of many reads (e.g. to examine repeat size and structure in an STR disease context), it is generally necessary to align the reads to a reference genome, filter to a region of interest, and clip long reads if they extend beyond the region of interest. Sections **3.2**–**3.3** provide detailed examples for using *minimap2* and *SAMtools* to process genome-wide reads down to a single curated FASTQ file per participant that can be parsed by `read_[modified_]fastq()`. However, this results in a mixture of forward and reverse reads relative to the region of interest. To enable consistent visualisation, *ggDNAvis* is able to intelligently reverse-complement only reverse reads by referencing a separate ‘metadata’ dataframe, which must contain columns for read ID and read direction.^1^ Metadata can be constructed manually for small datasets, or automatically using *SAMtools* (using the same aligned BAM file from which the input FASTQ is generated) per the code below:

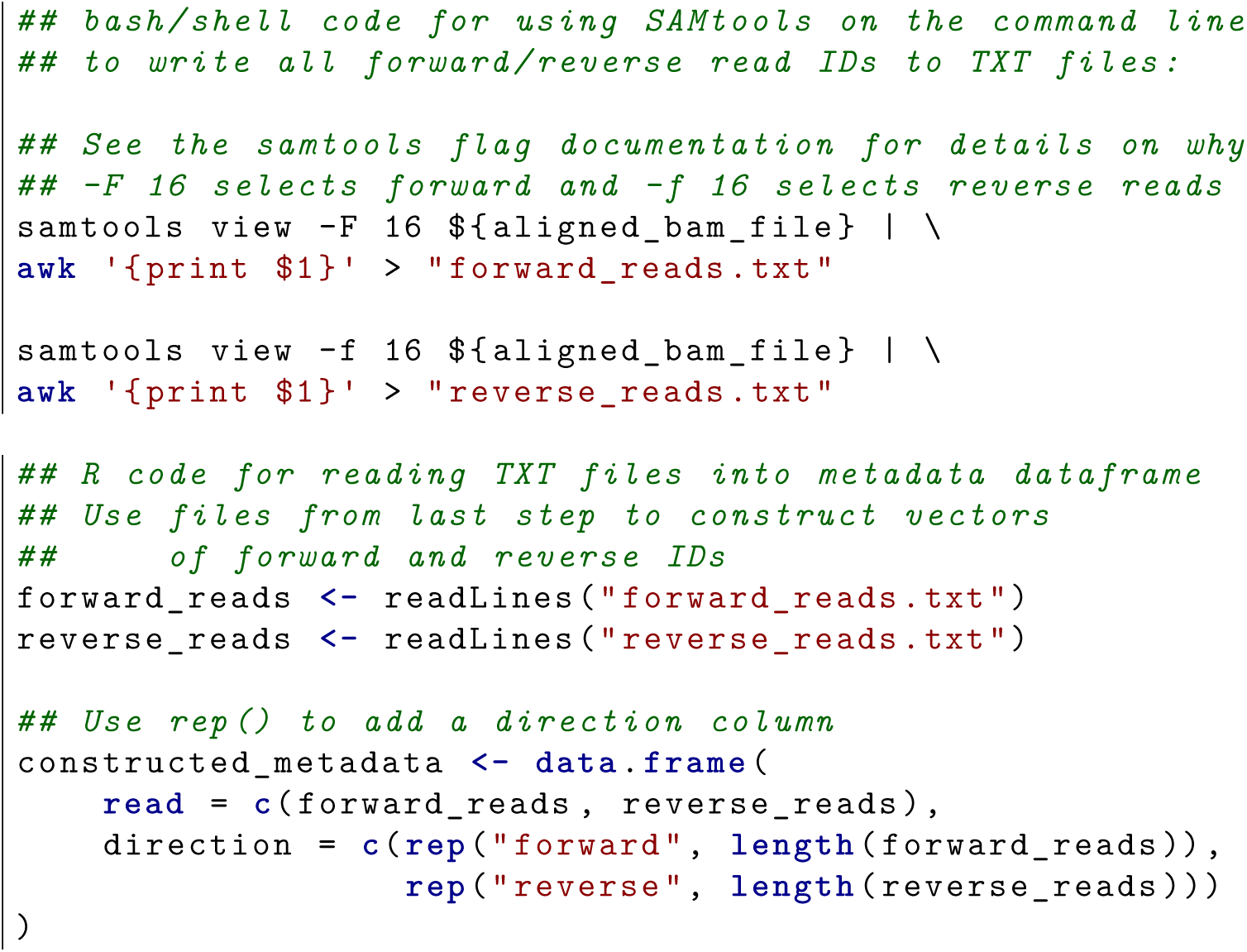

The metadata can then be merged with the FASTQ dataframe via merge_fastq_with_metadata(), which automatically uses the direction column to identify reverse reads. Reverse reads can be either preserved (**Figure 2a**), inverted (**Figure 2b**), or reverse-complemented (**Figure 2c–d**). If reverse-complemented, users can choose whether A maps to T or U to allow RNA compatibility. Likewise, metadata can be merged with modified FASTQ data via merge_methylation_with_metadata() to map the probabilities of various base modifications onto the visualisation. *ggDNAvis* maps modification information to original forward and reverse reads without recalculation (**Figure 2a**); to inverted reverse reads by reading modification probabilities in reverse order and recalculating modification positions relative to the start of the read (**Figure 2b**); and to reverse-complemented reverse reads by assigning modification probabilities either to the base complementary to that assessed (**Figure 2c**) or offsetting that by one base (**Figure 2d**). The latter is useful for modifications that occur once per palindromic motif, most commonly CG dinucleotides.

#### 2.1.3 Sequence extraction from dataframe

Once a FASTQ has been parsed and merged into a dataframe, *ggDNAvis* extracts values from the dataframe for visualisation. For single sequence visualisation, any character/string containing A/C/G/T/U is valid input,^2^ which can be created by hand or read from a particular cell of a FASTQ dataframe. This is then automatically split across multiple lines within the visualise_single_ sequence() function.

For multiple sequences/methylation visualisation, the order of the various reads should be arranged logically to communicate patterns in the data. The `extract_and_sort_sequences()` and `extract_ and_sort_methylation()` functions both read from the merged metadata-FASTQ dataframe to produce a vector of sequences (and, for methylation, vectors of modification locations and probabilities) for input to the ultimate visualise_<many_sequences/methylation>() function. In doing so, the grouping_levels argument allows the user to specify a hierarchical organisation of sequences, such as top-level grouping of reads by allele followed by second-level grouping by participant in a human disease genetics application (e.g. **Figure 5**). Grouping is implemented recursively: for each grouping variable, the data is successively subset to each value of the variable, then if the grouping variable is the lowest-level then the sequences are retrieved as-is. Conversely, if the grouping variable is not the lowest-level then sequences are retrieved via recursive function call to add spacing for lower-level variables. Length-0 strings (“”) for blank spacing lines are added after each set of sequences unless the value is the last of the variable.

### 2.2 Tiled rendering system

#### 2.2.1 Preprocessing

The high-level implementation approach of *ggDNAvis* is transforming the input DNA sequence/modification information into one or more *ggplot2*-readable dataframes, which are then plotted using the ‘tile’ geom.

The three major visualisation functions (of single sequences, multiple sequences, and methylation) each operate on character vectors where each element contains the information for one line. In multiple sequence visualisation, the input is a single character vector with one sequence/read per line. For methylation visualisation, the input is three character vectors containing sequence, modification location, and modification probability information respectively, again with one sequence/read per line. For single sequence visualisation, the input sequence is a length-1 character vector (i.e. one sequence), which needs to be split into a vector of substrings to be visualised across multiple lines. For a sequence of length *l* with user-specified line width *k*, line *i* is the substring from position *k*(*i* − 1) + 1 to *ki* while *ki* ≤ *l*. If *l* % *k >* 0, there is a final incomplete line from *k*(*i* − 1) + 1 to *l*. This vector of substrings is then equivalent to the vector input to multiple sequence or methylation visualisation, so downstream processing is done identically.

All three main visualisation functions allow annotation of the positions along each line. For single sequences, index annotations are on every line and are cumulative, while for multiple sequences/methylation they are on a user-specified subset of lines and reset each new line. This is implemented by the user passing an integer vector of the lines which should be annotated. Additional blank/spacer lines are then inserted immediately before or after each selected line (according to the index_ annotations_above Boolean argument), with the number of spacer lines inserted determined by ⌈*v*⌉ where *v* is the user-specified vertical alignment of the annotation text. In the special case of single sequence visualisation, every line of the split-sequence vector is annotated and the spacing between sequence lines is overridden from ⌈*v*⌉ to any user-specified spacing integer.

Ultimately, the result is vectors of sequences/locations/probabilities in the final arrangement from top to bottom, including any additional spacer lines added for index annotations. These vectors are then converted to numerical matrices to draw the coloured boxes, where 0–4 designate colours for sequence visualisation and −2–255 designate colours for methylation visualisation (see **2.2.2**). Furthermore, the vectors are read directly to rasterise x/y/value dataframes of text to be superimposed within each box.

#### 2.2.2 Rasterisation

Each *ggDNAvis* image is rendered as a grid occupying the coordinate space from (0, 0) to (1, 1). The grid is subdivided into *n* × *k* rectangles where *n* is the number of lines (sequence lines + spacer lines) and *k* is the maximum sequence length per line. Accordingly, each rectangle has a height of 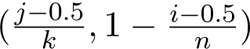 and a width of 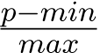. When exported to the final image file via ggplot2::ggsave(), the export width and height are set such that each rectangle becomes a 1-inch square, which becomes a *dpi* × *dpi* pixels square for any specified *dpi* value (controlled by the pixels_per_base argument).

To render the sequence lines as *n* × *k* rectangles, a family of rasterise_ functions takes a matrix (for the coloured boxes) or vectors of each line (for the superimposed text) organised from top to bottom, indexed such that *i* = 1 represents the top line and *j* = 1 represents the leftmost base of each line. The coordinates of the centre of each rectangle *i, j* are then 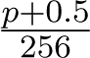. The rasterise_functions return a dataframe of these x and y coordinates for each base, along with a value column storing the value to be drawn.

For the coloured base boxes in single-/many-sequence visualisation, value is a numeric encoding of the base alphabetically from A = 1 to T/U = 4, or 0 for the background. For the coloured base boxes in methylation visualisation, value is the 0–255 integer probability of modification for assessed bases, −1 for non-assessed bases, or −2 for the background. For the sequence text inside the boxes, value is the letter directly. For the probability text inside methylation-visualisation boxes, value is the probability rounded to a specific precision and returned as a character value e.g. “256.00”. Prior to rounding, methylation probabilities are also scaled according to a user-specific min and max, with initial 0–255 integer probability *p* being transformed via 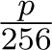. The default *min* = −0.5 and *max* = 256 scale the probability to 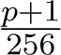 i.e. the centre of the 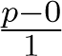 to 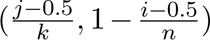 probability space (Bonfield & Marshall, 2024). Alternatively, setting *min* = 0 and *max* = 1 results in 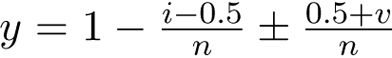 i.e. does not change the original 0–255 integer. Other scalings are possible as the parameters are simply numeric values, but produce a warning as they may be difficult to interpret.

Index annotations along each line are rasterised slightly differently, as they need to be positioned above or below each base rather than centred vertically within it. Relative to the centre of the annotated base rectangle at 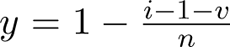, the index annotation is shifted half a rectangle up/down (to the vertical edge of the annotated rectangle) and a further *v* rectangle heights into the next i.e. 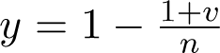, simplifying to 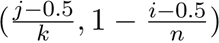 if the annotation is above and 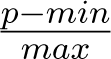 if the annotation is below.

#### 2.2.3 Appearance customisation

Finally, the rasterised dataframes are used as input for *ggplot2* to draw the visualisations. If box outlines, index annotations, or superimposed text within the boxes are required, geom_tile() is used to draw the boxes centred at 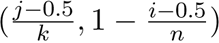 with width 1/*k* and height 1/*n*. In the simple case where only the coloured boxes are required, geom_raster() is used for an identical result with somewhat better performance.

Common appearance options are exposed as arguments, including colours, line widths, text sizes, margin etc. If further customisation is desired, the plot is returned as a ggplot object when return = TRUE is set, so further themes, geoms etc can be added just like any other ggplot. **Figure 2** is an example of a *ggDNAvis* visualisation being extended with additional ggplot layers. However, adding downstream geoms occurs after (thus is mutually exclusive with) built-in export via the filename argument, which uses the number and maximum character length of the sequence vector along with the margin setting to automatically set ggsave() width and height perfectly for each 1/*k* × 1/*n* rectangle to be exported as a *dpi* × *dpi* square. It is assumed that any users returning and further modifying the ggplot are experienced enough to be able to work out the correct dimensions for subsequent export.

For single-sequence and multiple-sequence visualisation, a key appearance choice is the colours used for each base. Six palette options are provided in the sequence_colour_palettes data, being:

- [A C G T] ggplot_style: the default 4-category colour palette in *ggplot2* of red (#F8766D), green (#7CAE00), cyan (#00BFC4), and purple (#C77CFF).
- [A C G T] bright_pale: pastel yellow (#FFDD00), green (#40C000), blue (#00A0FF), and red (#FF4E4E).
- [A C G T] bright_pale_2: pastel yellow (#FFDD00), slightly lighter green (#30EC00), blue (#00A0FF), and red (#FF4E4E).
- [A C G T] bright_deep: darker, richer tones of yellow/gold (#FFAA00), green (#00BC00), blue (#0000DC), and red (#FF1E1E) intended for use with white text.
- [A C G T] sanger: green (#00B200), blue (#0000FF), black (#000000), and red (#FF0000), inspired by Sanger sequencing chromatograms.
- [A C G T] accessible: light green (#B2DF8A), dark green (#33A02C), dark blue (#1F78B4), and light blue (#A6CEE3), suggested by colorbrewer2.org (Harrower & Brewer, 2003) as the only colourblind-friendly 4-category qualitative palette.

One important colour-customisation feature is the ability to ‘clamp’ the gradient of the modification probability scale in visualise_methylation(). By default, the low_colour is used for probability 0 and high_colour used for probability 255, with intermediate values interpolated by scale_fill_ gradient(). However, low_clamp and high_clamp values can be set such that low_colour is used for all probabilities ≤low_clamp, high_colour is used for all probabilities ≥high_clamp, and colour interpolation only occurs over the low_clamp to high_clamp range. Additionally, clamping can be used to make binary methylation calls, such as by setting low_clamp = 127 and high_clamp = 128 to classify all probabilities ≤127 as ‘unmethylated’ and all probabilities ≥128 as ‘methylated’ with no intermediate options (e.g. **Figure 5b**) — this results in the commonly used ‘lollipop’ style methylation plots (Rauch & Pfeifer, 2011), especially with white as low_colour and black as high_colour. Scale bars showing the exact colour scale including clamping can then be produced via visualise_ methylation_colour_scale() (e.g. **Figures 5–6**).

## 3 Results

A comprehensive explanation of all the possible *ggDNAvis* outputs is available from the *ggDNAvis* user guide. To briefly illustrate the three key output types available, publicly available data is used for three STR genes: *NOTCH2NLC* (NIID), *HTT* (HD), and *FMR1* (FXS).

### 3.1 Single sequence visualisation (*NOTCH2NLC*)

As discussed in **1**, the single sequence visualisation capabilities of *ggDNAvis* were designed to emulate the sequence visualisation style employed by Sone et al. (2019) (**Figure 1a**). In **Figure 3**, all four sequences shown in **Figure 1** were re-visualised using *ggDNAvis*’s visualise_single_sequence().

The F1-1 expanded *NOTCH2NLC* sequence from Sone et al. (2019) appears extremely similar to the original visualisation, but with index annotations spaced every 15 bases rather than every 25 to better align with the trinucleotide repeat units, and with the final base annotated at position 531 (**Figure 3a**).^3^ The F1-III-5 sequence from Yu, Luan et al. (2021) is now visualised as a regular grid rather than irregular Times New Roman text, and has positional information via the index annotations (**Figure 3b**). The Patient 14 Allele 2 sequence from Fitrah et al. (2023) is now visualised as a grid with consistent box sizes, line lengths, and line spacing and correct index annotations (**Figure 3c**). Finally, the F1-III-10 sequence from Yu, Deng et al. (2021) is also now a grid with correct numbering and length calculations (**Figure 3d**). This illustrates the utility of *ggDNAvis* to quickly and easily generate readable, customisable grid-based sequence visualisations with automatically and correctly calculated index (base position) annotations.

### 3.2 Multiple sequence visualisation (*HTT*)

To illustrate *ggDNAvis* multiple sequence visualisation, publicly available Cas-enriched Nanopore sequencing data from an HD cell line (Fang et al., 2022) was downloaded in FASTQ format from the Sequence Read Archive (sample ND33392, accession SRR13068459, from study SRP292765). Using *minimap2* v2.30-r1287 (H. Li, 2018), reads were mapped against Chromosome 4 of the GRCh38.p14 human reference genome (NCBI reference sequence NC_000004.12). Using *SAMtools* v1.23 (H. Li et al., 2009), reads were filtered to only those spanning the *HTT* repeat locus and hard-clipped to the CAG repeat plus 24 bp flanking sequence (6 bp upstream, 18 bp downstream) to ensure reliable insertion detection (clipped to Chr 4 positions 3,074,871–3,074,951, with the reference CAG repeat located at 3,074,877–3,074,933 followed by an additional CAACAG not counted in the repeat length per Walker, 2007).^4^ Forward and reverse read IDs were identified using the -F/-f 16 flag in samtools view and written to TXT via *awk* v20200816 (Aho et al., 1979), while the final clipped reads were written to FASTQ via samtools fastq. This mini-pipeline ran in under 2 minutes on a Macbook Air M1 8 Gb from many_sequences_htt.sh.

Reads were then loaded into *R* v4.5.2 via *ggDNAvis*’s read_fastq() and metadata was created based on the TXT lists of forward and reverse reads. These dataframes were merged via merge_fastq_ with_metadata(), using reverse_complement_mode = “reverse_only” to invert reverse reads to 3’–5’ (complementary to 5’–3’ forward reads). To display reads of the wildtype and expanded allele separately, alleles were defined via a clipped-read length threshold of 138 bp (see **Supplementary Figure S1**), equivalent to <u>^138*−*24^</u> = 38 CAG repeats. A random subsample of 250 reads was taken for visualisation, and extracted into a sequence vector via `extract_and_sort_sequences()` with top-level grouping by allele and second-level grouping via direction. This vector was then visualised via visualise_many_sequences() using the bright_deep palette, with additional geom_text markup in *ggplot2* v4.0.2, to create **Figure 4**. Full *R* code is available from many_sequences_htt.R.

In **Figure 4**, the bimodal distribution of *HTT* read lengths was apparent, with a large gap in repeat length between expanded and wildtype alleles and little intra-allele repeat length variation. The 5’-CAG-3’ (or 3’-GTC-5’ in reverse reads) repeat structure was clearly visible in all reads, with occasional substitutions or small insertions/deletions likely resulting from Nanopore sequencing errors. Interestingly, in several forward reads (both expanded and wildtype) there was a pattern of CAG being interrupted by CTT(GGC)*_n_* before returning to CAG. There was not an equivalent GAA(CCG)*_n_*interruption in reverse reads, indicating that this was most likely a recurring error in basecalling rather than a biological occurrence.

These kinds of observations, and many others given appropriate data, illustrate the utility of *ggDNAvis* for visually analysing and comparing multi-read sequencing data via easy-to-create, high-quality renders.

### 3.3 Methylation visualisation (*FMR1*)

To illustrate *ggDNAvis* methylation visualisation, raw Nanopore sequencing data of two XY male FXS cell lines (Coriell Institute NA04026 and NA05131), one XX female heterozygous *FMR1* premutation carrier (NA06905), and one XX female unaffected control (NA12878), sequenced by Stevanovski et al. (2022), was downloaded in BLOW5 format from the Sequence Read Archive (study SRP349335; files GBXM123343.tar, MBXM107326.tar, MBXM032249.tar, and MBXM044264.tar respectively) via download_fmr1.sh.

BLOW5 Nanopore signal files were converted to POD5 via *blue-crab* v0.5.0 (Gamaarachchi et al., 2022) then basecalled to BAM format in *dorado* v0.9.6+0949 (Oxford Nanopore, 2025) using the dna_r9.4.1_e8_sup@v3.3_5mCG_5hmCG@v0 maximum-quality CG hydroxy/methylation model. Using *SAMtools* v1.23 and *minimap2* v2.30-r1287 in parallel for each sample, reads were aligned against the GRCh38.p14 human reference genome (ENSEMBL Homo_sapiens.GRCh38.dna.primary_ assembly.fa.gz release 115), using the samtools fastq -T MM,ML and minimap2 -y options to retain modification information in FASTQ headers. Reads were then filtered to only those completely overlapping the *FMR1* repeat locus and hard-clipped to the CGG repeat plus 43 bp flanking sequence (19 bp upstream, 24 bp downstream) to ensure reliable insertion detection (clipped to Chr X positions 147,912,032–147,912,134, with the reference repeat located at 147,912,051–147,912,110). MM tags store modification locations relative to the original start of the read, so *modkit* v0.6.1 (Oxford Nanopore, 2026) was used to repair modification information in clipped reads. As in **3.2**, forward/reverse read ID lists were written to TXT and final clipped reads were written to FASTQ, with modification information copied to header rows via -T MM,ML. This pipeline was executed on the Research and Education Advanced Network New Zealand (REANNZ) high-performance computing platform via methylation_fmr1.sh.

For each cell line, reads were loaded into *R* via *ggDNAvis*’s read_modified_fastq() and metadata was created based on the TXT lists of forward and reverse reads. Methylation dataframes for all cell lines were then concatenated into a single dataframe, as were metadata dataframes.^5^ Methylation data and metadata were merged via merge_methylation_with_metadata(), using reverse_complement_ mode = “DNA” and reversed_location_offset = 1 to reverse-complement reverse reads and map C to C to identically match forward reads (see **Figure 2**). The total modification probability for each assessed C was determined by summing the integer scores of methylation and hydroxymethylation probability. *FMR1* repeats are typically categorised as wildtype below approximately 50 CGG units, premutation from 50–200 units, and full mutation above 200 units (Oostra & Willemsen, 2009) — reads were thus categorised based on clipped repeat length with ≤200 bp designated wildtype, 201–1,000 bp designated premutation, and >1,000 bp designated full mutation (see **Supplementary Figure S2**). Combined methylation probability for the full dataset was visualised grouped by allele then cell line in **Figure 5**, showcasing different options for colour scheme and probability clamping. Furthermore, methylation probabilities for NA12878 (unaffected XX control = homozygous wildtype) were visualised in **Figure 6**, showcasing different options for methylation text display. Full *R* code is available from methylation_fmr1.R.

In **Figure 5**, it was clear that the expanded (premutation and full mutation) *FMR1* reads were generally modified (methylated or hydroxymethylated), while wildtype reads were generally unmodified. **Figure 5a** showed that most CG sites had either very low modification probability (blue) or very high modification probability (red), with a few having intermediate probability (purple). **Figure 5b** showed that while the majority of reads followed the expanded = (hydroxy)methylated, wildtype = unmethylated pattern, there were some highly methylated wildtype reads, particularly of NA12878 (unaffected control).

There are three settings for sequence_text_type in visualise_methylation(): “none” (**Figure 5**), “sequence” (**Figure 6a**), and “probability” (**Figure 6b–c**). Sequence visualisation is self-explanatory and renders identically to visualise_many_sequences() but over boxes coloured by methylation probability instead of by base. Probability visualisation is based on the 0–255 integer probability scores — as discussed in **2.2.2**, any scaling 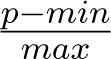 is possible, but the most useful options are sequence_text_scaling = c(−0.5, 256) to scale to 0–1 probabilities (**Figure 6b**) or c(0, 1) to leave as 0–255 integers (**Figure 6c**).

This extensive customisability allows *ggDNAvis* to provide flexible methods for visualising DNA modification information from modified FASTQ files and organising visualisations according to supplementary metadata.

## 4 Discussion

### 4.1 Advantages

#### 4.1.1 Easy generation of high-quality DNA visualisations

The core purpose of *ggDNAvis* is enabling authors to quickly and easily make high-quality renders of DNA information. The key design philosophy was allowing users to specify intentions such as spacing, colouring, or which lines of sequence to index-annotate at what frequency, while automatically managing the implementations thereof to reduce manual burden and therefore potential for errors. Comparing **Figure 1** and **Figure 3**, it is clear that features such as automatic layout and index calculation reduce the time, effort, and tedium of creating consensus sequence visualisations for authors — thereby also improving the clarity and numerical accuracy of the figures to make them easier to interpret for audiences. Between the three core visualisation functions of visualise_single_ sequence(), visualise_many_sequences(), and visualise_methylation(), a wide range of use cases are supported, from educators needing to display a custom sequence for usage in an exam to medical researchers compiling and comparing sequence and/or methylation data from Nanopore sequencing of many participants with STR expansions.

#### 4.1.2 Accessible but powerful

*ggDNAvis* was designed to be easy and approachable to get started, while being powerful and customisable for more advanced users. The *Shiny* web-app interface provides a quick-and-easy GUI for basic DNA visualisation even for users totally unfamiliar with *R* (**Figure 7**). While GUI interaction has limited reproducibility and automatability, care has been taken to implement saving and loading visualisation settings to reduce the need to manually re-configure each time the app is loaded. Furthermore, input is accepted either as text entry (straightforward) or file upload (faster for large datasets) to facilitate user agency in balancing simplicity against speed/power. Most configuration settings are accessible through the GUI, but menus are designed to hide irrelevant options (e.g. if sequence_text_type is set to “none”, then controls for probability scaling and rounding are hidden as they would have no effect) and provide sensible presets (e.g. the ‘probability’ and ‘integer’ modes seen in **Figure 6b–c**) so that less intuitive settings such as arbitrary numerical probability scaling are exposed only when ‘custom’ is explicitly chosen.

**Figure 7:**
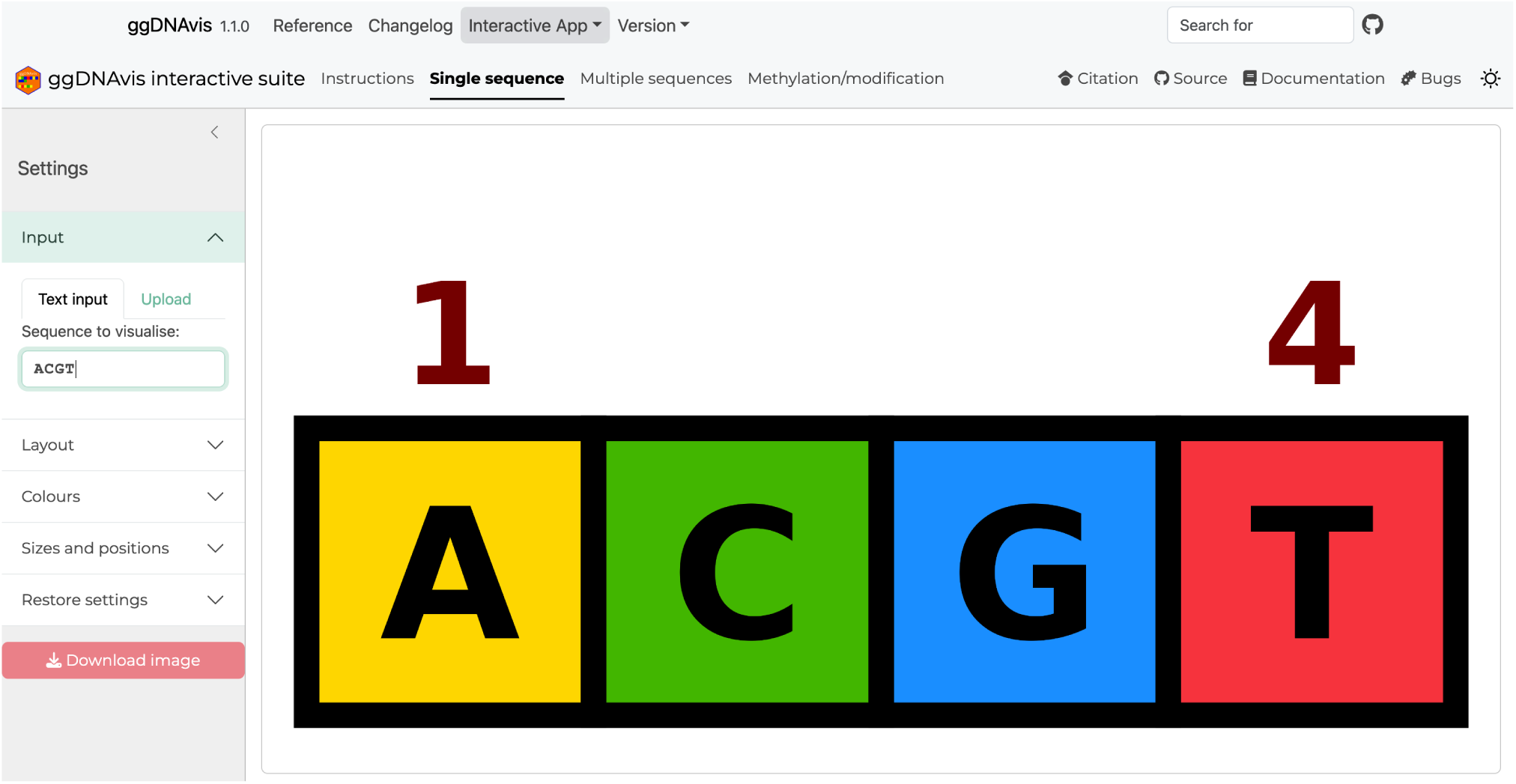
*Shiny* graphical user interface for *ggDNAvis*. The ‘Single sequence’ interface is currently selected, providing access to visualise_single_sequence(). Similar interfaces exist for visualise_many_sequences() and visualise_methylation(), accessed via the top menu bar. Sequence information can be input directly as text or uploaded as a FASTA/FASTQ file. Visualisation parameters can be controlled from the left menu bar, and can be saved to or restored from a JSON file. Final image can be saved to a PNG file via the ‘Download image’ button. Screenshot taken from *ggDNAvis* v1.1.0 *Shiny* interface on MacOS.

For intermediate users who want to use the package within *R*, the documentation website opens with a quickstart guide with succinct, complete examples of reading input files, processing data, and creating visualisations for each of the three visualisation types. Subsequently, comprehensive explanations are given of all major *ggDNAvis* functionality, organised to gradually build up complexity and customisability for each function. This complements the standard *R* package manual, which lists all functions in alphabetical order, detailing their descriptions, arguments, return values, and examples — an invaluable reference index, but not organised with ease of learning in mind.

Finally, advanced users are able to use *ggDNAvis* within the context of the broader *ggplot2* ecosystem because of the option to return as a ggplot object rather than exporting directly to PNG. While subsequent manual image export then does require precisely calculating target width and height (so is not recommended for beginners), the huge advantage is the ability to add any additional layers/geoms through *ggplot2*. For all *ggDNAvis* figures in this manuscript, this kind of post-hoc markup was employed to add titles, directions, and annotations. Furthermore, multiple visualisations can be composited directly within *R*, either by combining ggplot objects using packages such as *patchwork* (Pedersen, 2025) and *cowplot* (Wilke, 2025) (**Figure 3**) or by combining exported PNGs using *magick* (Ooms, 2026) (**Figures 4–6**). These advanced uses enable annotating and combining *ggDNAvis* visualisations into complex, publication-ready figures entirely within *R*, though of course there is the option to manipulate exported PNGs in other graphics software for users who are more comfortable doing so. The code to generate all *ggDNAvis*-constructed figures is freely available in the manuscript/directory of the source code repository as examples of extending *ggDNAvis* visualisations with other *ggplot2* ecosystem functionality.

#### 4.1.3 Reference genome-free visualisation

One particular benefit of *ggDNAvis* for the specific use-case of visualising STR expansions is that it operates on the reads themselves without requiring a reference genome. This means repeat expansions, which are necessarily ‘insertions’ relative to the reference genome, can be displayed in full and compared against each other rather than being relegated to an ‘insertion’ box against the reference. When *HTT* reads from the same sample used in **Figure 4** were viewed in *IGV* (**Figure 8**), the difference was immediately apparent: *IGV* displays the reference sequence and each read’s differences against it, while *ggDNAvis* shows each read’s sequence independently.

**Figure 8:**
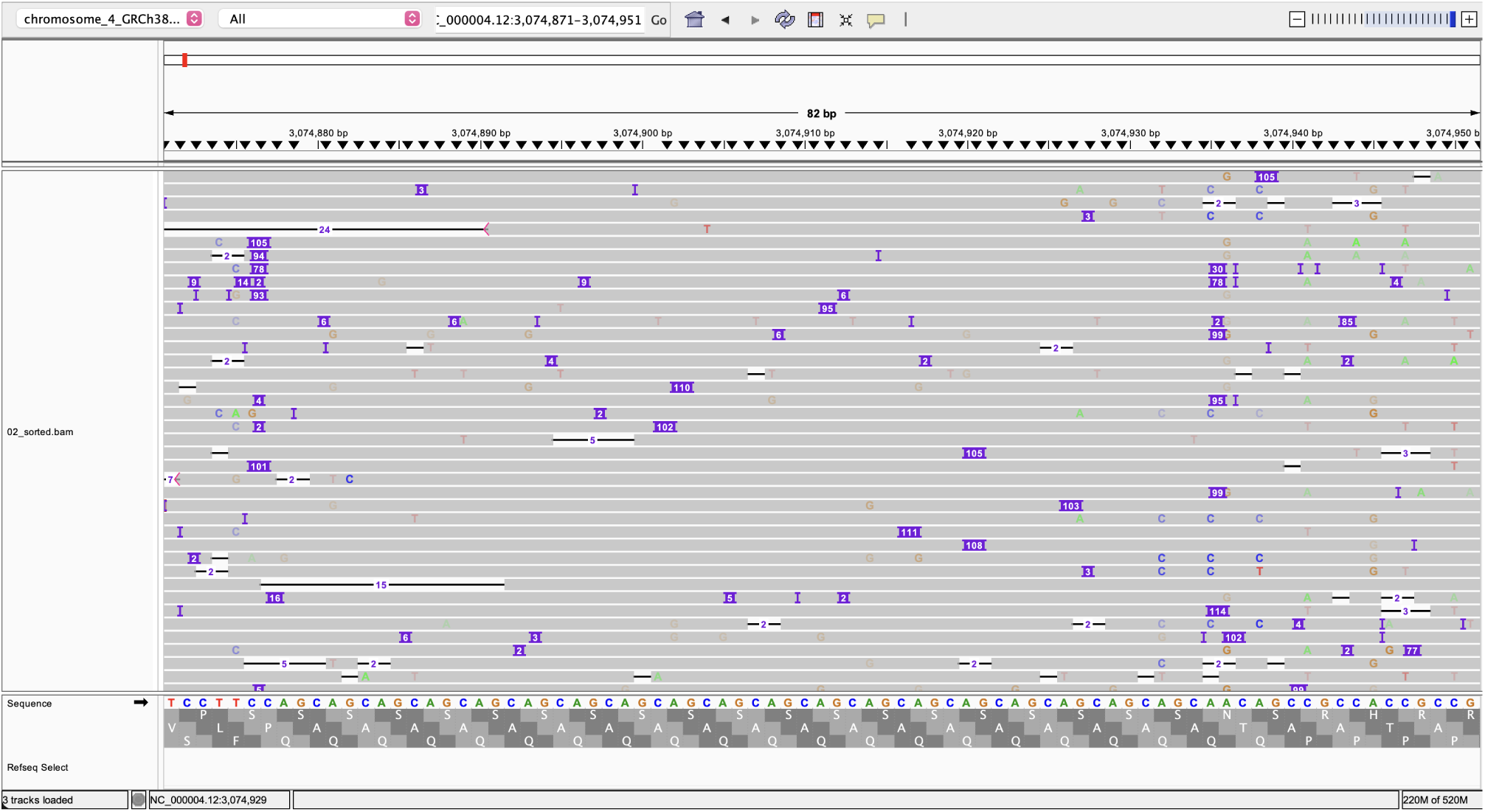
*HTT* reads from Nanopore DNA sequencing of HD fibroblasts visualised using *IGV*. Input data identical to Figure 4 i.e. sequencing data from Fang et al. (2022), downloaded from Sequence Read Archive (sample ND33392, accession SRR13068459, from study SRP292765), shown from Chr 4 positions 3,074,871–3,074,951. In *IGV*, expanded CAG repeats are shown as purple ‘insertions’ with the number of additional bases relative to reference in white text, but the expanded sequences themselves are not easily seen or compared. Data loaded from output/02_sorted.bam generated by many_sequences_htt.sh (see **3.2**). Screenshot taken from *IGV* v2.19.2 (Robinson et al., 2011) on MacOS.

*IGV* remains exceptionally useful for understanding the correspondence between the reads and the reference genome, and it was used heavily here when designing pipelines to extract and clip reads precisely over the repeat loci to use as *ggDNAvis* input. However, the specific use-case of visualising a full repeat sequence in many Nanopore reads is the exact purpose of visualise_many_sequences, and is therefore a particular strength of *ggDNAvis* compared to other visualisation software.

### 4.2 Limitations

#### 4.2.1 Performance

While effort has been taken to optimise performance where possible — for example, mathematical operations such as rasterisation (see **2.2.2**) are vectorised rather than done in nested for-loops — there are inherent limitations in the speed of rendering images with hundreds of squares and writing millions of pixels to disc. Therefore, the comparatively fast data processing operations involved in parsing FASTQ files and manipulating the sequences have been designed for simplicity rather than speed. For example, using list columns for modification locations and probabilities would avoid integer vector-character conversions via string_to_vector() and vector_to_string(), but the time savings would be negligible compared to saving images, while also increasing user-facing complexity of reading and writing *ggDNAvis* dataframes. Likewise, base *R* holds whole dataframes in memory, limiting the size of FASTQ that can be imported, but such immense datasets could not be visualised because, again, the visualisation export process is much more demanding than reading/parsing the datasets into memory. Overall, while there are inefficiencies in the implementation of some *ggDNAvis* processes within *R* (not to mention inefficiencies in *R* itself which could be sped up in a faster language), the inherent bottleneck in rendering and exporting the visualisations means further optimising data processing could not result in meaningfully faster start-to-finish runtimes.

#### 4.2.2 Square grid rendering system

The entire implementation of *ggDNAvis* relies on using geom_tile() (or occasionally geom_raster()) from *ggplot2* to draw grids of identically sized squares. This means one major customisability limitation is that line spacing (whether the gaps between line sections of sequence in visualise_single_ sequence() or allowable group separation values in extract_and_sort_<sequences/methylation>()) must be integer. Allowing non-integer spacing would be a welcome customisation option, but would require major changes to the rendering system and associated calculations (e.g. export px dimensions) that rely on the assumption of 1 base/spacer = 1 line. There are no current plans to enact these changes.

#### 4.2.3 Preprocessing requirements

*ggDNAvis* is designed to be useful in a wide range of contexts and purposes. The most basic functionality of visualising a single DNA/RNA molecule is indeed near-universal as it can accept any DNA sequence from any source. However, the more complex multiple sequence visualisation is most suitable for long-read sequencing, and generally requires preprocessing to isolate reads from a single locus of interest. While there is no technical limitation preventing FASTQ files from short-read (e.g. Illumina) sequencing from being loaded and visualised in *ggDNAvis*, the ∼150 bp reads would appear messy and inconsistent because they would not be aligned and *ggDNAvis* shows reads in absolute terms rather than relative to a reference genome — for short reads, *IGV* would generally be more appropriate. Conversely, long reads (often tens or hundreds of kilobases in length) can fully span a locus of interest and can be clipped to the coordinates of interest such that every clipped read has identical start and stop locations (point mutations or basecalling errors notwithstanding), allowing useful comparisons when visualised together. This requires some experience in using bioinformatics software such as *minimap2* and *SAMtools* to perform the read alignment, filtering, and clipping, raising the barrier to entry for *ggDNAvis* multiple sequence visualisation for users with little bioinformatics experience.

Methylation/modification visualisation, meanwhile, absolutely requires sequence and modification information for individual DNA molecules i.e. amplification-free modification-capable sequencing. In most cases, this means recent Oxford Nanopore sequencing with signal-level (FAST5/POD5/SLOW5) data available, or BAM files which have already been basecalled with a modification-capable model. As with multiple sequence visualisation, preprocessing is virtually mandatory to produce useful input files, including filtering reads to a region of interest, clipping to precisely the target locus, and an additional step of repairing the modification information after clipping.

Therefore, single sequence visualisation is straightforward and universal, while multi-read visualisation (of sequence or modification information) requires specific input data and preprocessing steps. Nonetheless, for the subset of geneticists with appropriate long-read sequencing data (e.g. STR disease researchers), *ggDNAvis* multi-read visualisation offers a useful approach distinct from other visualisation software.

### 4.3 Conclusions

Overall, *ggDNAvis* offers easy-to-use software for creating highly customisable DNA visualisations of a single sequence, multiple sequences, or methylation of multiple Nanopore reads. The *Shiny* web interface provides accessibility and a low barrier to entry for quick visualisation projects, while integration of the *R* package with the broader *ggplot2* ecosystem allows advanced users to create complex annotations and composite visualisations. Detailed documentation and worked examples in the documentation website facilitate learning and understanding the various features. By providing a fast and reliable way to generate DNA visualisations, we hope to empower authors and improve the quality of visualisations while reducing the difficulty of making them.

## Declarations

### Ethics approval and consent to participate

Not applicable — all data used in this study was previously published and is publicly available from the referenced manuscripts and repositories.

### Consent for publication

Not applicable — all data used in this study was previously published and is publicly available from the referenced manuscripts and repositories.

### Availability of data and materials

- Project name: *ggDNAvis*
- **Project home page:** https://ejade42.github.io/ggDNAvis/
- **Operating system(s):** MacOS, Windows, Linux
- **Programming language:** *R*
- **Other requirements:** *ggplot2* v4.0.0 or higher
- License: MIT
- Any restrictions to use by non-academics: License needed

All source code for *ggDNAvis* is available under MIT license from the source code repository. All code used to process example data and generate figures is available from the manuscript/directory of the source code repository. All data used is publicly available from the Sequence Read Archive, specifically studies SRP292765 (Fang et al., 2022) and SRP349335 (Stevanovski et al., 2022).

Full links for all data and materials used in this study:

|  |  |
| --- | --- |
| *ggDNAvis* on CRAN | https://cran.r-project.org/package=ggDNAvis |
| Source code repository | https://github.com/ejade42/ggDNAvis |
| Code for figures | https://github.com/ejade42/ggDNAvis/blob/v1.1.0/manuscript |
| Documentation website | https://ejade42.github.io/ggDNAvis/ |
| Interactive *Shiny* app | https://ejade42.github.io/ggDNAvis/articles/interactive_app.html |
| Fang et al. (2022) | https://trace.ncbi.nlm.nih.gov/Traces/?view=study&acc=SRP292765 |
| Stevanovski et al. (2022) | https://trace.ncbi.nlm.nih.gov/Traces/?view=study&acc=SRP349335 |

## Generative AI usage

LLMs (*ChatGPT* v4o and *Gemini* v3 Pro) were used to assist with specific function implementations in *R*, *LATEX*, and *Bash*. All generated code was thoroughly human-checked. Generative AI was not used in any capacity for writing the text of this manuscript.

## Authors’ contributions

**E.J.** (corresponding author): Conceptualization, Data Curation, Investigation, Methodology, Project Administration, Software, Visualization, Writing – Original Draft. **E.L.S.**: Funding Acquisition, Project Administration, Resources, Supervision, Writing – Review & Editing.

## Acknowledgments

This work was funded by the Neurological Foundation of New Zealand, the Health Research Council of New Zealand, and the associates of the late Marcus Gerbich. We thank Jessie Jacobsen and Klaus Lehnert for their assistance proof-reading and fact-checking this manuscript. We acknowledge the use of Research and Education Advanced Network New Zealand (REANNZ), formerly New Zealand eScience Infrastructure (NeSI), high-performance computing facilities. We thank Susanna Palmer, Olivia Brighouse, Elani Richards, and Emily Caldelari-Hume for their contributions to testing *ggDNAvis*.

## Footnotes

1 As of *ggDNAvis* v1.1.0, reverse reads are ‘forward-ified’ in this manner, but additionally forward reads are ‘reverseified’. This is important for genes on the minus strand, where the local ‘forward’ direction is ‘reverse’ relative to the global reference genome.

2 Note that base *R* character values are intentionally used rather than Biostrings from the *Bioconductor* ecosystem (Gentleman et al., 2004), so that *ggDNAvis* can be easily installed via install.packages() without needing to install and manage *Bioconductor* dependencies.

3 These minor tweaks are fully customisable so the Sone et al. (2019) figure could be faithfully reproduced by setting index_annotation_interval = 25 and index_annotation_always_last_base = FALSE if desired.

4 Flanking sequences were required because *minimap2* sometimes places the ‘insertions’ (i.e. repeat expansions) slightly outside the reference repeat locus coordinates, in which case they get clipped away unless sufficient flanking sequence is also included.

5 One read (9fd72b80-b4ca-4070-8058-5de62cbb2b64) of NA05131 was excluded because the read mapping, prior to manual hard-clipping, was soft-clipped 2,354 bp left but did not have an ‘insertion’ — this suggests the read ended midway through the repeat expansion or soon enough into the unique flanking sequence that *minimap2* was unable to correctly map the expansion.

