## Supplementary material for "*ggDNAvis*: a *ggplot2*-based *R* package for creating high-quality DNA sequence and modification visualisations"

27th October 2026

**References** **7**

### List of Figures

S1 Histogram of *HTT* repeat lengths justifying allele separation threshold at 138 bp. . . 5
S2 Histogram of *FMR1* repeat lengths justifying allele separation thresholds at 200 bp and 1,000 bp. . . . . 6

### 1 Detailed FASTQ parsing information

#### 1.1 Standard FASTQ parsing

Second-generation sequencing-via-synthesis platforms such as Illumina provide information on the sequence of each read (Ambardar et al., 2016), which is easily stored in standard FASTQ (Cock et al., 2010). The FASTQ format is plain text with 4 lines per entry: read identifier/header, sequence, spacer, quality. Therefore, lines are read into *R* as a character vector (one value per line), then divided by type according to line number modulo 4. Vectors of read ID, sequence, and quality (subset from the full FASTQ-lines vector via line number modulo 4 == 1, 2, and 0 respectively) are used to construct a dataframe, with a `sequence_length` column optionally constructed by counting the number of characters in each element of the `sequence` column. As writing to FASTQ via *SAMtools* tends to leave '@' symbols at the start of each read, `read_fastq()` and `read_modified_fastq()` also have a `strip_at` Boolean argument to remove one leading '@' from each read ID that begins with '@'.

#### 1.2 Modified FASTQ parsing

##### 1.2.1 `MM` and `ML` tags explanation

Unlike second-generation sequencing-by-synthesis, third-generation Nanopore long-read sequencing operates by threading DNA through a nanoscopic protein pore and detecting changes in electrical signal characteristic of each base passing through (Ambardar et al., 2016). When base modifications (notably 5-methylcytosine) are present, such as on natural DNA extracted from cells (not PCR-amplified), such modifications result in distinct electrical signal changes compared to unmodified bases which can be identified during basecalling (Simpson et al., 2017). Basecalling software such as Oxford Nanopore Technologies' *Dorado* (Oxford Nanopore, 2025) frequently outputs to SAM/BAM format (Li et al., 2009), where base modification information is stored in the `MM` and `ML` tags.

According to the *SAMtools* tags specifications (Bonfield & Marshall, 2024), `MM` stores the *locations* at which modifications of a particular type were assessed (e.g. methylation of cytosines of `CG` dinucleotides). It starts with an `MM:Z:` identifier, followed by a modification code and comma-separated location list for each modification type, with different modification types separated by semicolons. The modification code is a short string describing the type of information encoded, such as `C+m?` for methylation of cytosine bases.[1] The comma-separated location list then stores the number of bases of that type (e.g. cytosines) skipped before each assessed base (e.g. `C`s of `CG`s). For example, `C+m?,2,5,0,3` encodes that the first two cytosines were not assessed for methylation, the third was, the next five were not, the ninth was, the tenth was, the next three were not, and the fourteenth was. This could be expressed as `00100000110001`, where the `0`s represent `C`s that were not assessed for methylation and `1`s represents `C`s that were assessed (e.g. `C`s of `CG`s). In this representation, the numbers in the `MM` tag correspond to the number of `0`s before each `1`. A complete `MM` tag might therefore look like `MM:Z:C+h?,2,5,0,3;C+m?,2,5,0,3`—this tag would indicate that both hydroxymethylation and methylation have been assessed for the third, ninth, tenth, and fourteenth `C`s in the read.

Conversely, the `ML` tag encodes the *probability* of modification for each site at which modification was assessed. Like `MM`, it starts with a `ML:B:C:` identifier, followed by a comma then a comma-separated list of 8-bit integers (`0` to `255`). These integers encode probabilities, with integer $p$ covering the probability space from $\frac{p}{256}$ to $\frac{p+1}{256}$. For example, a score of `100` encodes that the probability of modification for that base was between $\frac{100}{256} = 0.391$ and $\frac{101}{256} = 0.395$. Thus, the centre of the probability space is $\frac{p+0.5}{256}$ e.g. $\frac{100.5}{256} = 0.393$ for a score of `100`. There is one `ML` number for each `MM` number regardless of modification type, with the first portion of the `ML` list representing probabilities for the locations listed in the first modification type within `MM`, and then the second portion of the `ML` list representing probabilities for the second modification type within `ML` and so on. Therefore, the `MM:Z:C+h?,2,5,0,3;C+m?,2,5,0,3` tag from above would be accompanied by an 8-number `ML` sequence, say `MM:B:C,200,25,0,75,20,100,150,125`. This would be interpreted as the third `C` having

[1] The '`?`' specifies that no information is given about the likelihood of modification for skipped bases, whereas a '`.`' would indicate that skipped bases have low modification probability.

approximately a $\frac{200.5}{256} = 0.783$ probability of hydroxymethylation and $\frac{20.5}{256} = 0.080$ of methylation, the ninth `C` having approximately $\frac{25.5}{256} = 0.100$ probability of hydroxymethylation and $\frac{100.5}{256} = 0.393$ of methylation, and so on. These are mutually exclusive, as the fifth carbon of the cytosine cannot be simultaneously methylated and hydroxymethylated, so the remaining probability is the chance of being unmodified (e.g. approximately $\frac{(255-200-20)+0.5}{256} = \frac{35.5}{256} = 0.139$ for the third `C`).

##### 1.2.2 `MM` and `ML` tags parsing

`MM` and `ML` tags natively store modification information within a SAM/BAM file. However, *SAMtools* also has the functionality to copy tags (tab-separated) into the header lines of a FASTQ via `samtools fastq -T MM,ML input.bam > output.fastq`. *ggDNAvis* can then parse these into a dataframe via `read_modified_fastq()`. The sequence and quality lines are exactly the same as normal FASTQ and are parsed identically, but the header row is now split by tab to separate the read ID from the `MM` and `ML` tags.

The `MM` (location) tags are separated by modification type and converted from storing the number of skipped target bases between each assessed base to simply storing the absolute index (starting at 1) of each assessed base. For example, in the sequence `TACTCGACACG` there are 4 `C`s at indices 3, 5, 8, and 10, with the second and fourth `C` being part of a `CG` dinucleotide. Therefore, the `MM` tag might include `C+m?,1,1` to indicate methylation assessment of the second and fourth `C`s. *ggDNAvis* would then take the cumulative sum along the skips vector with every element increased by 1 (e.g. `cumsum(c(1, 1) + 1)`) to produce a vector of which `C`s were assessed (e.g. `c(2, 4)`). This vector is then used to subset a vector of all `C` locations (e.g. `c(3, 5, 8, 10)[c(2, 4)]`) to produce a final vector of all assessed base locations (e.g. `c(5, 10)`). This is calculated for each modification type of each read in a functional programming paradigm via `lapply()` and `sapply()`.

The `ML` (probability) tag for each read is parsed by taking a subset of the single long comma-separated list based on the length of the skips vector for each modification type. For example, if the first modification type has skips `1,0,2` (length 3) and the second modification type has skips `0,5` (length 2), then elements 1–3 of the probability vector would be the modification probabilities for the first modification type, and elements 4–5 would be the probabilities for the second modification type.

Ultimately, a dataframe is produced containing read ID, quality, sequence, and sequence length (as with parsing unmodified FASTQ), but with additional comma-separated-list columns of modification types present in each read and, for each modification type, locations and probabilities of each assessed base.

### 2 Example visualisation methodological details

#### 2.1 Multiple sequence visualisation (*HTT*) — allele separation

As mentioned in the main text, *HTT* wildtype and expanded alleles were separated by length with a threshold at 138 bp (38 `CAG`). The threshold for full-penetrance *HTT* pathogenicity is 41 `CAG` repeats, with reduced-penetrance pathogenicity from 36–40 repeats (Walker, 2007). However, the overwhelming majority of reads could be unambiguously assigned an allele regardless of precise threshold placement, with only 7 of the 3,701 filtered reads (0.19%) within ±5 bp inclusive of 138 bp and only 18 (0.49%) between 36 and 41 repeats (132–147 bp, inclusive). Therefore, the precise allele separation threshold was not important, as it would only affect the allele categorisation of a tiny minority of reads (**Supplementary Figure S1**). 138 bp was chosen because it corresponds to 38 `CAG` in the middle of the reduced-penetrance range and some specific value was needed, but any similar value could have been used with negligible impact on the outcomes.

#### 2.2 Methylation visualisation (*FMR1*) — allele separation

Likewise, *FMR1* wildtype, premutation, and full-mutation were separated by length with thresholds at 200 bp and 1,000 bp. This corresponded to trinucleotide lengths of 52.33 and 319 repeats, assuming exactly 43 bp flanking sequence. However, as Stevanovski et al. (2022) did not use Cas enrichment,

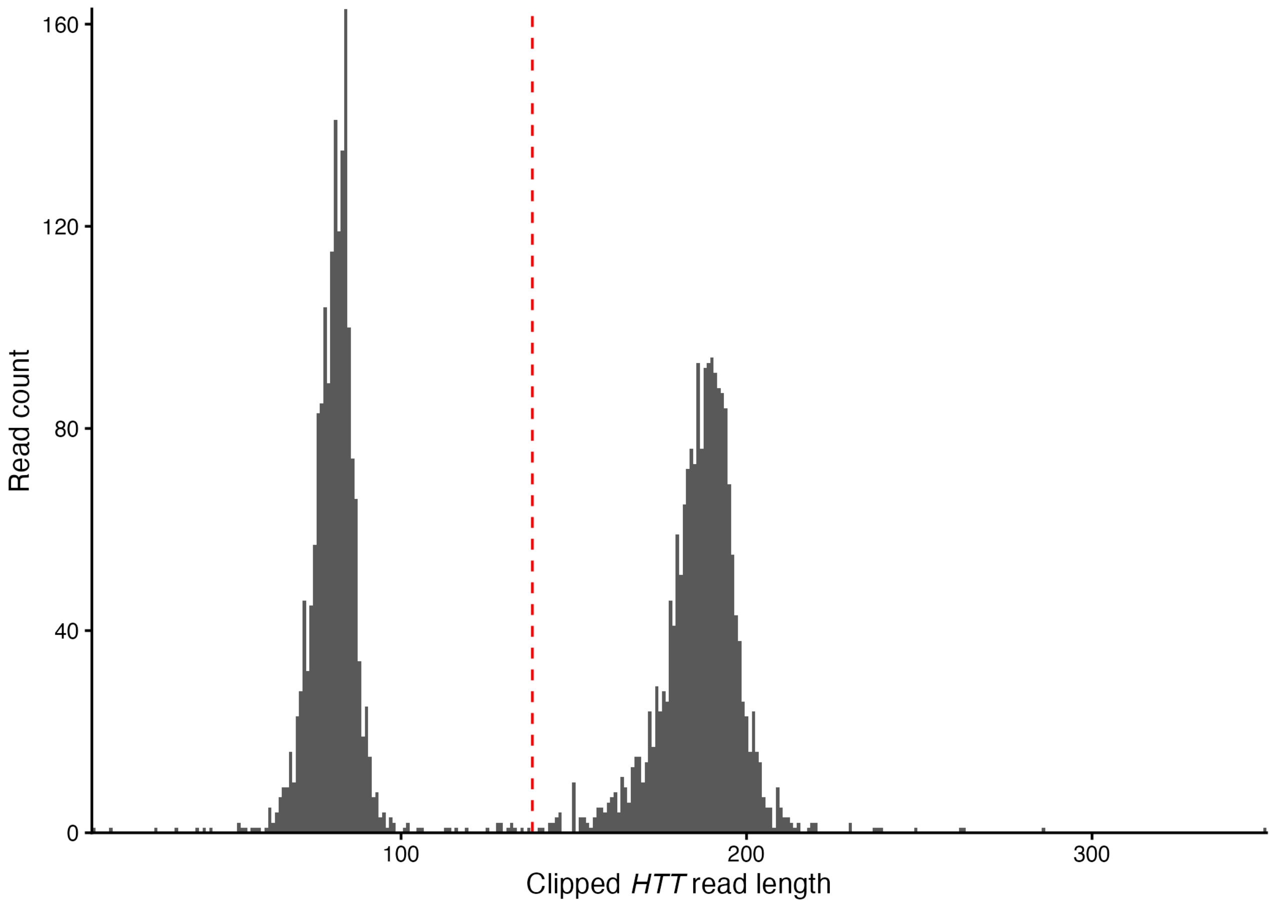

**Supplementary Figure S1: Histogram of *HTT* repeat lengths justifying wildtype/expanded allele separation threshold at 138 bp** (vertical red line). Repeat lengths were clearly bimodal and any threshold from approximately 100–150 bp would separate the distinct allele types. Sequencing data from Fang et al. (2022), downloaded from Sequence Read Archive (sample ND33392, accession SRR13068459, from study SRP292765). Reads were mapped, filtered, and clipped by `many_sequences_htt.sh`, then further processed and visualised by `many_sequences_htt.R`.

the *FMR1* coverage was low enough that there were zero reads of intermediate/ambiguous length — wildtype reads in this data were entirely from 19–32 repeats, premutation reads were 71.67–84.33, and full mutation reads were 406–724.67 (**Supplementary Figure S2**). Thus, any thresholds from 33–71 and 85–405 repeats would be unambiguous for this data, consistent with biological thresholds at 50 and 200 repeats in the literature (Oostra & Willemsen, 2009). The choices of 200 bp and 1,000 bp were purely for round numbers and could be moved anywhere within the gaps between allele categories with zero effect on allele categorisations.

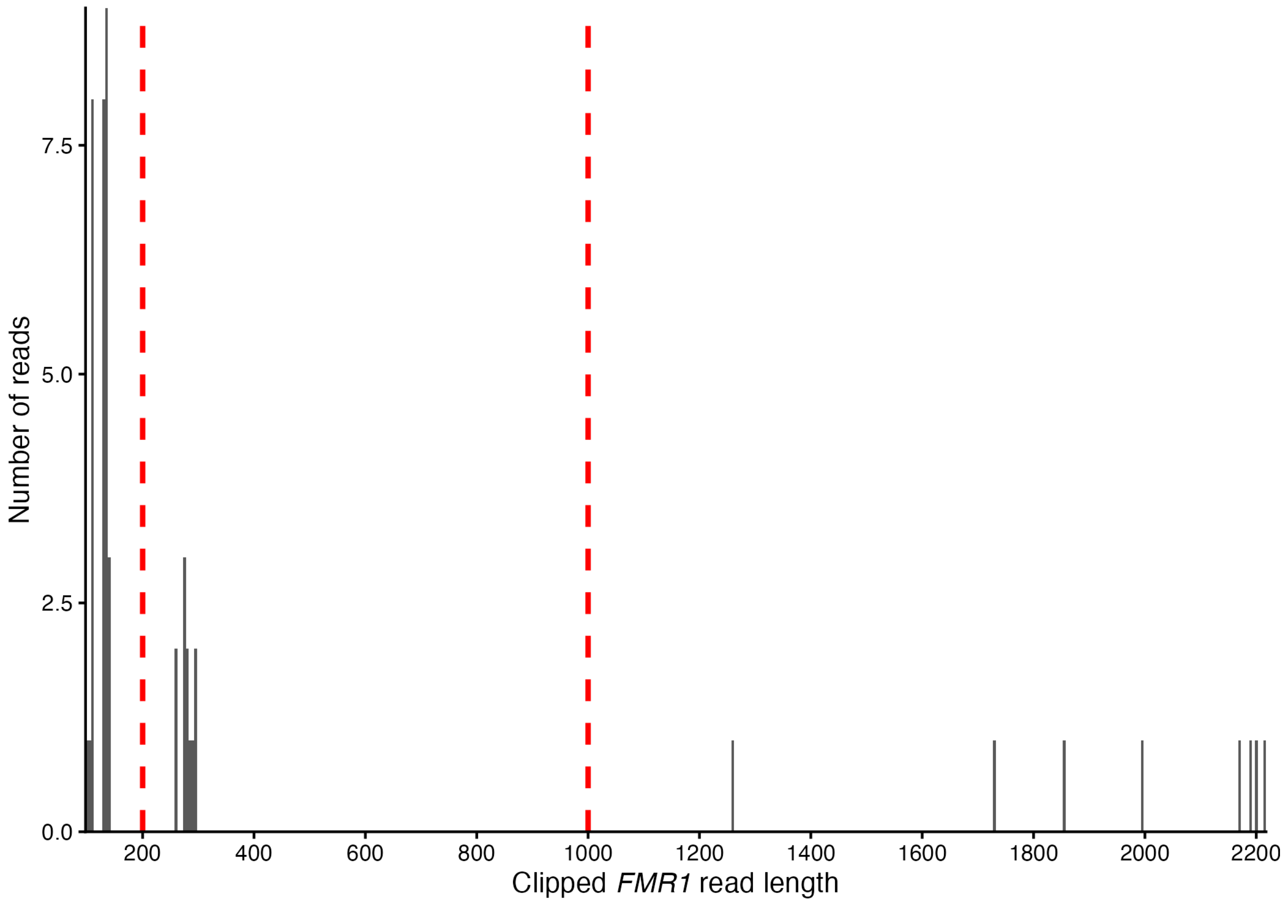

**Supplementary Figure S2: Histogram of *FMR1* repeat lengths justifying wildtype/premutation/full mutation allele separation thresholds at 200 bp and 1,000 bp** (vertical red lines). Repeat lengths were clearly trimodal and any threshold from approximately 150–250 bp and 300–1,250 bp would unambiguously separate the distinct allele types. Sequencing performed by Stevanovski et al. (2022) on Coriell Institute cell lines NA04026 (FXS XY male, hemizygous full mutation), NA05131 (FXS XY male, hemizygous full mutation), NA06905 (unaffected XX female, heterozygous premutation), and NA12878 (unaffected XX female, homozygous wildtype). Data downloaded from SRA study SRP349335 via `download_fmr1.sh`, processed via `methylation_fmr1.sh`, then figure generated by `methylation_fmr1.R`.
